# Resistance from Afar: Distal Mutation V36M Allosterically Modulates the Active Site to Accentuate Drug Resistance in HCV NS3/4A Protease

**DOI:** 10.1101/452284

**Authors:** Ayşegül Özen, Kuan-Hung Lin, Keith P Romano, Davide Tavella, Alicia Newton, Christos J. Petropoulos, Wei Huang, Cihan Aydin, Celia A. Schiffer

## Abstract

Hepatitis C virus rapidly evolves, conferring resistance to direct acting antivirals. While resistance via active site mutations in the viral NS3/4A protease has been well characterized, the mechanism for resistance of non-active site mutations is unclear. R155K and V36M often co-evolve and while R155K alters the electrostatic network at the binding site, V36M is more than 13 Å away. In this study the mechanism by which V36M confers resistance, in the context of R155K, is elucidated with drug susceptibility assays, crystal structures, and molecular dynamics (MD) simulations for three protease inhibitors: telaprevir, boceprevir and danoprevir. The R155K and R155K/V36M crystal structures differ in the α-2 helix and E2 strand near the active site, with alternative conformations at M36 and side chains of active site residues D168 and R123, revealing an allosteric coupling, which persists dynamically in MD simulations, between the distal mutation and the active site. This allosteric modulation validates the network hypothesis and elucidates how distal mutations confer resistance through propagation of conformational changes to the active site.

## Introduction

Hepatitis C (HCV) infection causes inflammation of the liver and can lead to cirrhosis and liver cancer if not treated effectively. HCV NS3/4A protease inhibitors (PIs) were the first class of direct-acting anti-virals (DAAs) approved by the FDA to be used in combination with peginterferon-alfa and ribavirin. These PIs included the covalent, acyclic inhibitors (telaprevir (Perni, Almquist et al. 2006, Kwong, Kauffman et al. 2011), boceprevir (Malcolm, Liu et al. 2006)) and a non-covalent, macrocyclic inhibitor (simeprevir (Raboisson, de Kock et al. 2008, Rosenquist, Samuelsson et al. 2014). Subsequently, macrocyclic PIs have been developed as second-generation DAAs, including danoprevir, grazoprevir, and recently FDA-approved glecaprevir.

PIs target the virally encoded serine protease NS3/4A. NS3 is a 631 amino-acid bifunctional protein, with a serine protease domain in the N-terminus followed by an NTPase/RNA helicase domain. The crosstalk between the two domains has been previously reported (Frick, Rypma et al. 2004, Beran and Pyle 2008, Beran, Lindenbach et al. 2009, Aydin, Mukherjee et al. 2013), but domains fold into active proteins independently. NS3/4A protease domain, a chymotrypsin-like fold with two β-barrel domains, has a catalytic triad (H57, D81, S139) located in a cleft separating the two barrels. NS4A, a 54-amino acid peptide, is required as a cofactor for optimal proteolytic activity. The central 11 amino acids of the cofactor inserts as a β-strand to the N-terminal β-barrel of NS3 to form the active enzyme (Yao, Reichert et al. 1999).

While PIs offer a subpopulation of patients improved sustained virologic response and the potential for shorter therapy duration (Jacobson, McHutchison et al. 2011, Poordad, McCone et al. 2011, Sherman, Flamm et al. 2011, Gane, Pockros et al. 2012), the efficacy of approved PIs is challenged by drug resistance (Halfon and Locarnini 2011, Halfon and Sarrazin 2012, Svarovskaia, Martin et al. 2012, Vermehren and Sarrazin 2012, Welsch and Zeuzem 2012). Mutations arise in the NS3/4A protease depending on the therapeutic regime; A156 mutates in response to treatment with linear ketoamide protease inhibitors (Kieffer, Sarrazin et al. 2007, Sarrazin, Kieffer et al. 2007, Susser, Welsch et al. 2009, Halfon and Sarrazin 2012, Svarovskaia, Martin et al. 2012, Vermehren and Sarrazin 2012, Welsch and Zeuzem 2012) while macrocyclic PIs mainly select for D168A and R155K variants (Manns, Reesink et al. 2011, Manns, Bourliere et al. 2011, Lim, Qin et al. 2012). Mutations at V36, T54, V36/R155 were initially reported to be associated with resistance to ketoamide inhibitors (Kieffer, Sarrazin et al. 2007, Sarrazin, Kieffer et al. 2007, Susser, Welsch et al. 2009). V36M/R155K mutations were observed in long-term follow-up of patients receiving boceprevir (Howe, Long et al. 2015). An interferon-free mericitabine/danoprevir combination therapy resulted in a confirmed viral breakthrough in 21% of patients (Gane, Pockros et al. 2012) and these patients had danoprevir-resistant variants, including V36M/R155K. Although the newer PIs have improved potency and cure rates in combination therapy, dual and triple mutant variants including those with V36M mutations still emerge in patients failing therapy (Sorbo, Cento et al. 2018)

We have previously elucidated the structural basis of resistance due to the active site mutation R155K by crystal structures (Romano, Ali et al. 2010, Romano, Ali et al. 2012). R155 participates in an electrostatic network of interactions along the substrate-binding surface, which involves residues D81-R155-D168-R123. This strong electrostatic network is associated with tighter substrate binding (Romano, Laine et al. 2011), therefore is critical in stabilization of the bound ligands. In fact, most early inhibitors, particularly the compounds with aromatic P2 moieties, pack against R155. Our crystal structures (Romano, Ali et al. 2010, Romano, Ali et al. 2012, Soumana, Ali et al. 2014) showed that the R155K mutation disrupts this network by eliminating one of the two hydrogen bonds of R155. As a result, the altered charge distribution along the binding surface affects the P2 cyclopentylproline and P4 cyclohexylalanine moieties of telaprevir whereas danoprevir loses a favorable cation-π interaction at the P2 isoindoline (Romano, Ali et al. 2012). MD simulations based on these crystal structures suggest that the destruction of the salt bridge between 168 and 155 may cause additional conformational changes in the binding pocket (Pan, Xue et al. 2012).

Unlike R155K, the molecular mechanism of resistance for V36M mutation is unknown. V36M is a mutation distal to the active site, located on A1 strand, > 13 Å away from the catalytic histidine (H57) and 15 Å from R155 (Figure 1). In the absence of crystal structures, molecular modeling led to the hypothesis that the distal mutations at V36 and T54 can impair interaction with telaprevir’s cyclopropyl group (Welsch, Domingues et al. 2008). However, no experimental data exists on the structural changes due to V36M and how the effects of this distal mutation may affect PI binding at the active site. We have previously developed the “network hypothesis” postulating that the effect of distal mutations in HIV-1 protease are propagated to the active site through a network of intra-molecular interactions to confer drug resistance (Ragland, Nalivaika et al. 2014). The network hypothesis may be applicable to distal mutations in HCV NS3/4A protease as well, if this is a general mechanism by which resistance occurs.

**Figure 1.**
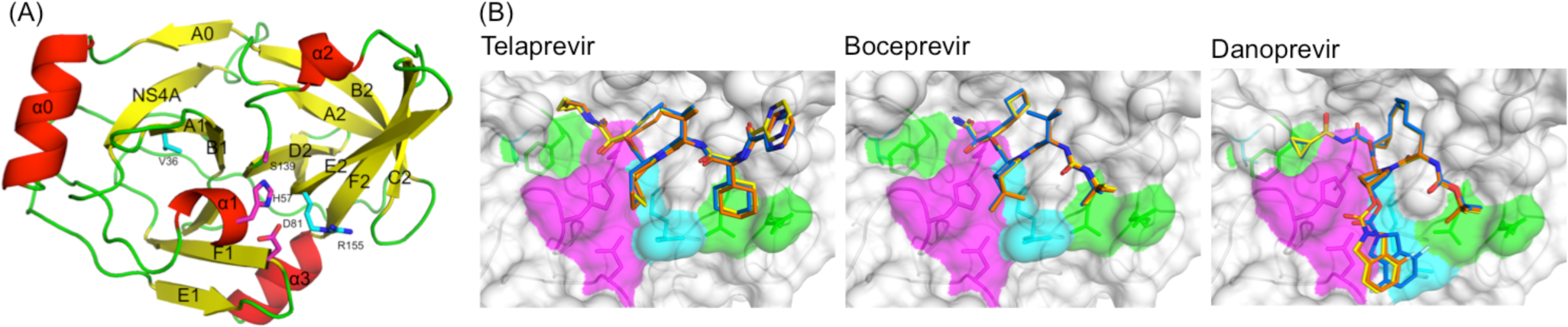
(A) Structure of HCV NS3/4A protease where α-helices, strands, and loops are colored in red, yellow, and green, respectively. Side chains of the catalytic triad are in magenta and the mutation sites R155K and V36 are in cyan. (B) Crystallographic binding modes of telaprevir, boceprevir, and danoprevir in the wild-type, R155K and R155K/V36M protease complexes. Protease and inhibitors are represented as surface and sticks, respectively. Side chains of key residues are also shown as sticks; the drug resistance mutation sites R155K and V36M (cyan), the catalytic triad H57-D81-S139 (magenta), and other binding site residues (green). Inhibitors in complex with wild-type, R155K and R155K/V36M proteases are shown in blue, orange and yellow respectively.

To understand why the distal mutation V36M is selected together with the active site mutation R155K to confer drug resistance, we determined the crystal structures of R155K/V36M double mutant HIV NS3/4A protease with three PIs: telaprevir, boceprevir, and danoprevir (Table S1). Through a combination of drug susceptibility assays (enzymatic and viral replicon), extensive structural analysis and MD simulations, we show that V36M accentuates resistance to PIs by allosterically modulating the conformation and dynamics of the active site. Most notably, the change in the shape of the binding site in NS3/4A protease is induced through a conformational shift in B1 strand. These findings provide key insights into the interplay between the structural and mutational plasticity of the enzyme in conferring drug resistance; validating the network hypothesis (Ragland, Nalivaika et al. 2014) and explaining how distal mutations are able to contribute to conferring drug resistance through alteration of the dynamic network and modulation of the active conformations.

## Results

To understand the molecular basis of the selection of V36M distal mutation under the pressure of PI including regimens, wild-type (WT) and resistant protease variants carrying R155K and R155K/V36M mutations were compared for binding telaprevir, boceprevir, and danoprevir. Crystal structures of the R155K/V36M double mutant protease were determined in complex with the three PIs (Table S1), and analyzed in comparison with our previously published crystal structures of the telaprevir- and danoprevir-bound R155K single mutant (Romano, Ali et al. 2012). For additional comparison, high-resolution crystal structures of apo and boceprevir-bound WT were also determined (Table S1) and carefully generated models of apo R155K and R155K/V36M variants (see Materials and Methods). These twelve (10 crystal and 2 model) resulting structures and their subsequent MD simulations (100 ns in triplicate) were analyzed for changes in protein structure, dynamics, and molecular interactions at the binding site. In addition, the susceptibility of the single and double mutant variants to the three PIs was determined both with enzymatic and viral assays.

### V36M Further Decreases Susceptibility of R155K variants to PIs

The activity of the three PIs against R155 and R155/V36M variants was assessed by enzymatic (Ki) and the cellular half-maximal (IC50) inhibition constants (Table 1). The enzyme inhibition constant against the full-length NS3/4A versus the protease domain alone was comparable, regardless of the PI or mutation introduced, suggesting a minimal role for the helicase domain in protease inhibitor binding. All three PIs are significantly less active against the double mutant than against the WT and R155K variants. Compared to WT enzyme, boceprevir has 7- and 31-fold less inhibitory activity against the R155K and R155K/V36M protease respectively (Table 1) while telaprevir loses even more activity, 24-fold against R155K and more than 200-fold against R155K/V36M. For danoprevir, the fold-reduction in Ki relative to WT is remarkably severe for both single and double mutants, 158-fold and 295-fold respectively. Despite this severe fold-change in potency, danoprevir is still the most potent among the three PIs against the resistant protease variants, due to an order-of-magnitude better potency against WT protease (1.2 nM versus 34.7 and 40.9 nM). While the double mutant has varying degrees of susceptibility to the three PIs, in all cases, V36M enhances resistance against both linear and macrocyclic compounds compared to R155K alone.

**Table 1.** Drug susceptibilities against wild-type and resistant HCV clones and inhibitory activities against NS3/4A proteases.

| **Full-length NS3/4A Enzyme - $K_i$ (nM)<sup>a</sup>** |  |  |  |
| --- | --- | --- | --- |
|  | ***WT*** | ***R155K*** | ***R155K/V36M*** |
| ***Telaprevir*** | 40.9 ± 3.7 | 824.0 ± 75.1 (20) | >10,000 (>244) |
| ***Boceprevir*** | 34.7 ± 2.9 | 390.8 ± 43.0 (11) | 1018.0 ± 192.3 (29) |
| ***Danoprevir*** | 1.2 ± 0.1 | 132.0 ± 18.0 (111) | 292.9 ± 38.6 (246) |
| **Protease domain $K_i$ (nM)<sup>a</sup>** |  |  |  |
|  | ***WT*** | ***R155K*** | ***R155K/V36M*** |
| ***Telaprevir*** | 33.3 ± 4.0 | 803.7 ± 89.9 (24) | 7342 ± 1281 (220) |
| ***Boceprevir*** | 35.4 ± 3.3 | 236.7 ± 44.3 (7) | 1097.0 ± 120.4 (31) |
| ***Danoprevir*** | 1.0 ± 0.1 | 157.9 ± 20.5 (158) | 295.5 ± 34.3 (295) |
| **Replicon $IC_{50}$ (nM)<sup>a</sup>** |  |  |  |
|  | ***WT*** | ***R155K*** | ***R155K/V36M*** |
| ***Telaprevir*** | 1349 | 4740 (3.5) | 15759 (12) |
| ***Boceprevir*** | 971 | 2788 (3) | 3941 (4) |
| ***Danoprevir*** | 0.24 | >100 (>416) | >100 (>416) |

<sup>a</sup>Numbers in parentheses reflect fold-change relative to wild-type; > indicates $IC_{50}$ and $K_i$ values higher than the maximum drug concentration tested in the assay.

Replicon-based cellular inhibition results correlate well with the enzyme inhibition constants. For all three PIs, the loss of antiviral activity (fold-change in IC50 relative to wild-type) against the R155K/V36M clones is substantial compared to the HCV clones carrying the R155K single mutation. In replicon assays, telaprevir and boceprevir lost more than 3-fold activity with R155K mutation while these linear compounds were 12- and 4-fold less potent against the R155K/V36M virus relative to wild-type. The IC50 of danoprevir was beyond the 100 nM assays limit, which corresponds to more than 416-fold loss of antiviral activity. In conclusion, the non-active site mutation V36M reduces the activity of all three PIs both on molecular and cellular levels in the presence of the active site mutation R155K.

### R155K/V36M Mutations Destabilize Danoprevir’s Large P2 Moiety

The overall binding modes of PIs were conserved regardless of the resistance mutations in the crystal structures (Figure 2). However, protease–inhibitor interactions were altered due to the mutations to varying extends. Danoprevir exhibits the largest shift due to the resistance mutations in the crystal structures. P2-isoindoline of danoprevir is in direct contact with the site of mutation, forming cation-π stacking interactions with R155, while the P2 groups of telaprevir and boceprevir are smaller and relatively distant to the 155 sidechain. The R155K mutation disrupts the favorable cation-π stacking and causes a shift in the danoprevir P2 moiety in the crystal structures of both the single and double mutant complexes (Figure 2 and 3A). These changes have been described as the mechanism underlying resistance of R155K protease to danoprevir (Romano, Ali et al. 2012). In addition to these structural changes, our MD simulations here reveal that the interactions of danoprevir’s P2 moiety with the protease are destabilized in both protease variants, with a considerably enhanced conformational flexibility of the bound inhibitor (Figure 3B). This enhanced conformational flexibility causes loss of favorable interactions even more than those observed in the crystal structures, including with the catalytic residues D81 and H57 (Figure 4 versus S1). The P2 moiety samples alternate conformations different than that in the starting crystal structure or the WT complex during the simulations and loses more than 6 kcal/mol in van der Waals interactions with the protease, mainly with residues 155–157 (Figure 5). In both single and double mutant variants, danoprevir also loses favorable packing of the P1’ moiety with F43 on the B1 strand (Figure 5), which interfaces both the active site and A1 strand containing the site of distal mutation. In the R155K/V36M protease complex, danoprevir forms new interactions with V158, S159, and T160 not present in WT and R155K variants (Figure 5). Previously we found these residues form more favorable interactions with the substrates than inhibitors (Ozen, Sherman et al. 2013). Thus, while the loss of van der Waals interactions in the danoprevir crystal structures are less pronounced and comparable for the single and double mutant variants, the destabilization and further loss of P2 interactions revealed in MD simulations correlate with the high fold-change in potency (Table 1).

**Figure 2.**
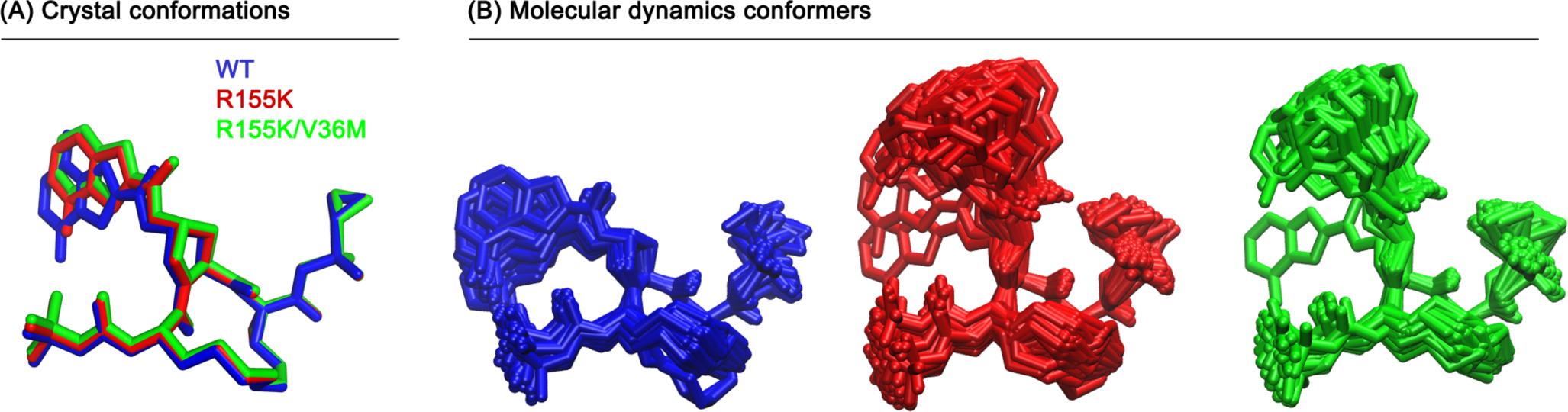
Conformational flexibility of danoprevir’s P2 moiety is enhanced by the molecular interactions destabilized due to R155K mutation. (A) Crystal structures and (B) representative snapshots from MD simulations with equal intervals of 6 ns superimposed on the C2, N2, C4, N5, C11, and C15 atoms of danoprevir.

**Figure 3.**
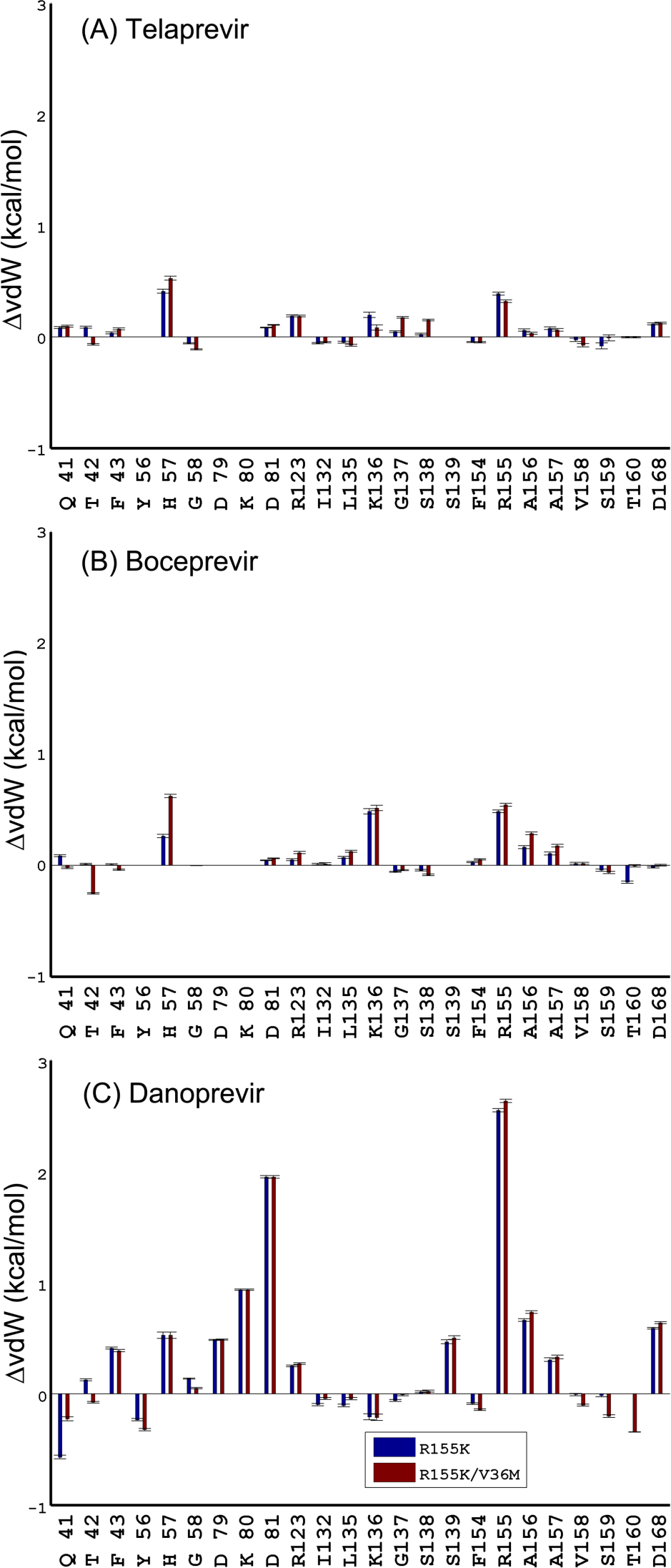
Changes in the van der Waals interactions of R155K and R155K/V36M protease residues relative to the wild-type protease for the complexes of (A) telaprevir, (B) boceprevir, (C) danoprevir from MD simulations.

**Figure 4.**
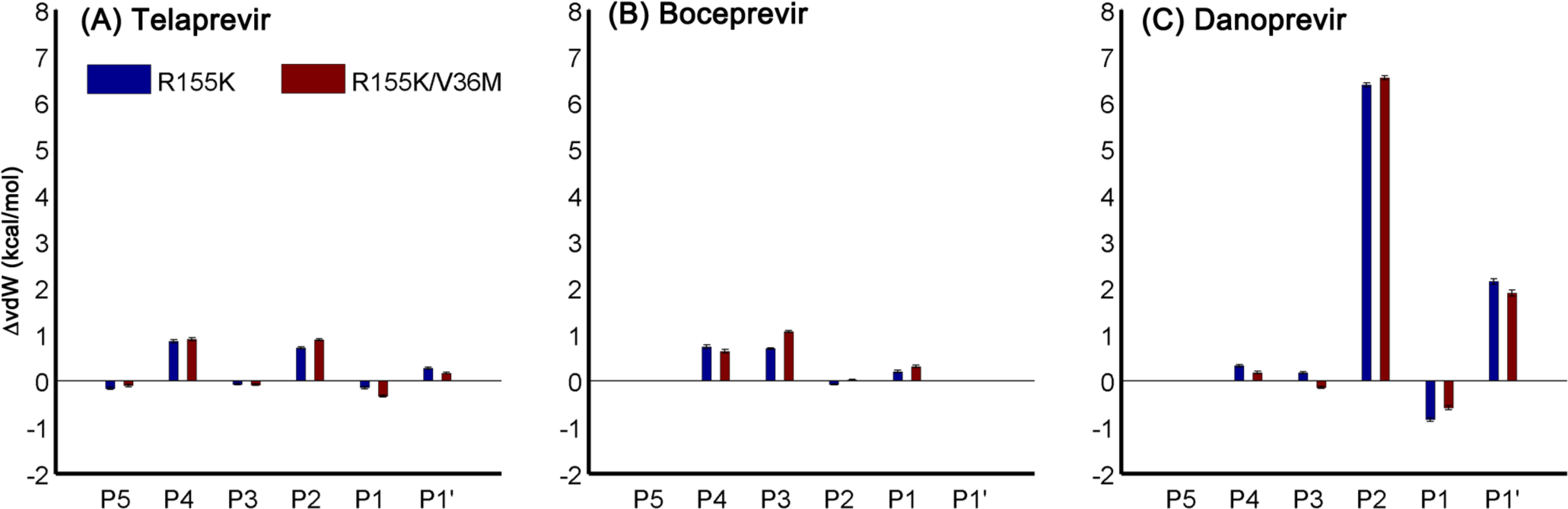
Changes in van der Waals interactions of (A) telaprevir, (B) boceprevir, and (C) danoprevir chemical moieties with the R155K and R155K/V36M proteases from MD simulations relative to the wild-type protease.

**Figure 5.**
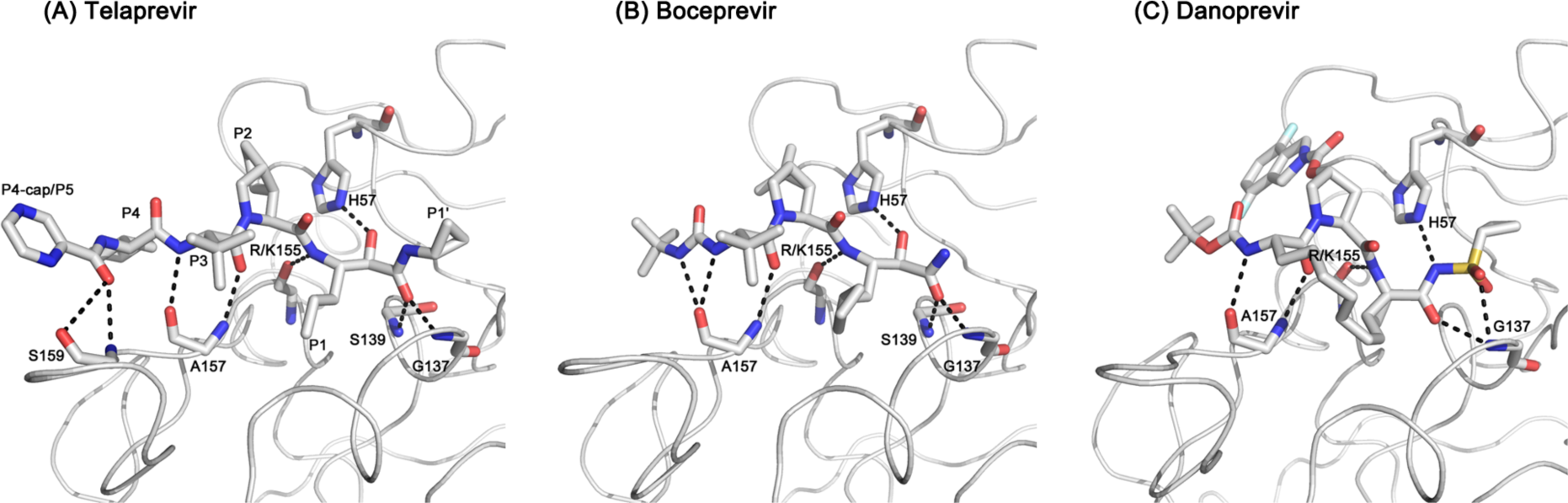
Protease-inhibitor hydrogen bonds in co-crystal structures. (A) Telaprevir, (B) boceprevir, (C) danoprevir. Hydrogen bonds are preserved in complexes of resistant variants; thus they are shown only in wild-type structures. See Table S2 for the resistant variants.

### R155K/V36M Mutations Impact Dynamic Packing of Linear Inhibitors at the Active Site

The two linear inhibitors with smaller P2 moieties, telaprevir and boceprevir, have comparable potencies against WT protease, but telaprevir is more susceptible to resistance due to the R155K and V36M mutations (Table 1). The loss of vdW interactions with the mutation site in R155K variants is much less pronounced in the two linear inhibitors compared to danoprevir, in both crystal structures and MD simulations (less than 1 kcal/mol, Figures 4 and S1), likely due to lack of a large P2 moiety packing against residue 155. Telaprevir loses favorable vdW interactions mainly with residue 155 and the catalytic H57 at its P2 and P4 positions, while boceprevir loses interactions also with K136, A156 and A157 (Figure 4).

In the crystal structures, the distal site mutation V36M causes further loss of vdW contacts with K136 and R155 for telaprevir and boceprevir, respectively (Figure S1), compared to R155K alone. However, comparison of the vdW energies in the crystal structures versus dynamic conformations sampled during MD indicates that the V36M distal mutation significantly alters the dynamic interactions between the inhibitors and the protease (Figures 4 and S1). Unlike in the crystal structures, the overall impact of resistance mutations is not localized to a single residue but dispersed through subtle changes in the vdW contacts over the binding surface altering the packing of inhibitors to the resistant variant proteases.

Key protease residue side chains in contact with telaprevir are perturbed by the R155K/V36M double mutation. The telaprevir-bound R155K/V36M protease crystal structure has multiple side chain conformations at the M36 position. Substitution of a valine at a relatively buried position to a larger methionine side chain alters the local packing, and the M36 side chain assumes these alternate conformations. This perturbation of side chain packing due to V36M mutation is transmitted to the protease active site causing two residues, D168 and R123, which are part of the active site electrostatic network, to have alternate conformations as well. Hence, in the double mutant, the active site electrostatic network is destabilized both due to active site R155K and the distal V36M mutations (Table 2). In addition, K136, which plays a role in substrate recognition through favorable electrostatic interactions with the acidic P6 position in the substrates, also makes favorable vdW contacts with telaprevir. However, no clear density was observed for the K136 sidechain in R155K/V36M co-crystal structure of telaprevir, suggesting that the distal V36M mutation has perturbed the interactions of K136 in α1 helix with telaprevir. Dramatic loss of affinity with R155K/V36M compared to the single mutant R155K may be partially due to these conformational changes observed in the crystal structures both in the binding site and at the distal site of mutation.

**Table 2.** Intramolecular salt bridges forming a network at the NS3/4A active site surface in crystal structures, and stabilities assessed by MD simulations.

|  |  | APO |  |  | Telaprevir |  |  | Boceprevir |  |  | Danoprevir |  |  |
| --- | --- | --- | --- | --- | --- | --- | --- | --- | --- | --- | --- | --- | --- |
|  |  | WT | R155K | R155K<br>V36M | WT | R155K | R155K<br>V36M | WT | R155K | R155K<br>V36M | WT | R155K | R155K<br>V36M |
| *Crystal Structures (Distance between the side chain oxygen and nitrogen in Å)* |  |  |  |  |  |  |  |  |  |  |  |  |  |
| H57 | D81 | 3.9 |  |  | 3.9 | 3.8 | 3.8 | 3.9 | 3.9 | 3.9 | 3.9 | 3.9 | 3.9 |
| D81 | R155 | 6.4 |  |  | 4.5 | - | - | 4.5 | 3.7 | 3.5 | - | - | - |
| R155 | D168 | 4.8 |  |  | 4.8 | 4.8 | 5.2 | 4.8 | - | - | 3.7 | - | - |
| D168 | R123 | 3.6 |  |  | 4.8 | 4.8 | 4.3 | 4.6 | 3.5 | 3.6 | 5.1 | 3.5 | 3.5 |
| D25 | R11 | 4.5 |  |  | 4.5 | 4.4 | 4.4 | 4.5 | 4.5 | 4.4 | 4.4 | 4.4 | 4.4 |
| E173 | E118 | 5.7 |  |  | 5.7 | 5.6 | 5.5 | 5.5 | 5.6 | 5.5 | 5.4 | 5.4 | 5.4 |
| E32 | R92<sup>a</sup> | 3.5 |  |  | - | - | - | 4.1 | - | 3.3 | - | - | - |
| E30 | R997 | 4.5 |  |  | 4.7 | 4.6 | 4.6 | 4.8 | 4.7 | 4.5 | 4.9 | 4.7 | 4.8 |
| D103 | R117 | 4.5 |  |  | 4.6 | 4.5 | 4.5 | 4.6 | 4.6 | 4.6 | 4.6 | 4.6 | 4.7 |
| D121<sup>b</sup> | R118<sup>b</sup> | - |  |  | - | - | - | - | - | - | 6.7 | - | - |
| D112<sup>b</sup> | R130<sup>b</sup> | - |  |  | - | - | - | - | - | - | - | 6.8 | 6.5 |
| *MD Simulations (% of the time the salt bridge criterion was satisfied)* |  |  |  |  |  |  |  |  |  |  |  |  |  |
| H57 | D81 | 100 | 99.4 | 99.9 | 93.9 | 29.1 | 34.6 | 100 | 100 | 100 | 100 | 22.7 | 30.5 |
| D81 | R155 | 10.3 | 12.2 | 16.4 | 95.8 | 89.3 | 83.3 | 85.5 | 2.6 | 3.9 | 0 | 83.9 | 78.4 |
| R155 | D168 | 98.3 | 5.6 | 3.1 | 86.2 | 23.8 | 23.9 | 79.6 | 65.0 | 67.3 | 100 | 22.2 | 21.3 |
| D168 | R123 | 21.7 | 96.2 | 99.7 | 88.5 | 96.0 | 95.1 | 88.7 | 91.7 | 84.9 | 22.3 | 69.5 | 90.8 |

<sup>a</sup>In only three structures, unambiguous electron density was observed for R92.
<sup>b</sup>All structures had relatively high B-factors for these residues.

### Resistance Mutations Cause Changes in the Active Site Electrostatic Network

Hydrogen bonds are significant contributors to inhibitor binding. The PIs make hydrogen bonds mainly with the backbone donors/acceptors in the binding site (Figure 6). As a result, the impact of R155K on the hydrogen-bonding network is minimal. Distance between the hydrogen bond donor and acceptor atoms vary 0.2 Å at most, staying the same for the most part in crystal structures. However in MD simulations, because of the structural rearrangement in the binding site caused by V36M, the likelihood of a hydrogen bond with the backbone oxygen decreases ∼20% for telaprevir and ∼10% for boceprevir (Table S2). The most striking difference between the crystal structures and MD simulations is that both telaprevir and danoprevir lose a hydrogen bond with the catalytic H57, which is observed ∼60% of the time between boceprevir and WT or R155K protease, but drops to 34% in R155K/V36M complex. In contrast, hydrogen bonds with R/K155 and A157 (on the E2 strand) are preserved in the MD simulations, suggesting that these bonds are key contributors to inhibitor binding. Hence, drug resistance mutations modulate the relative stability of certain inter-molecular hydrogen bonds in the conformational ensembles sampled in the MD simulations while the hydrogen bonds that are likely indispensable for binding remain stable across crystal structures and throughout the simulations of mutant complexes.

**Figure 6.**
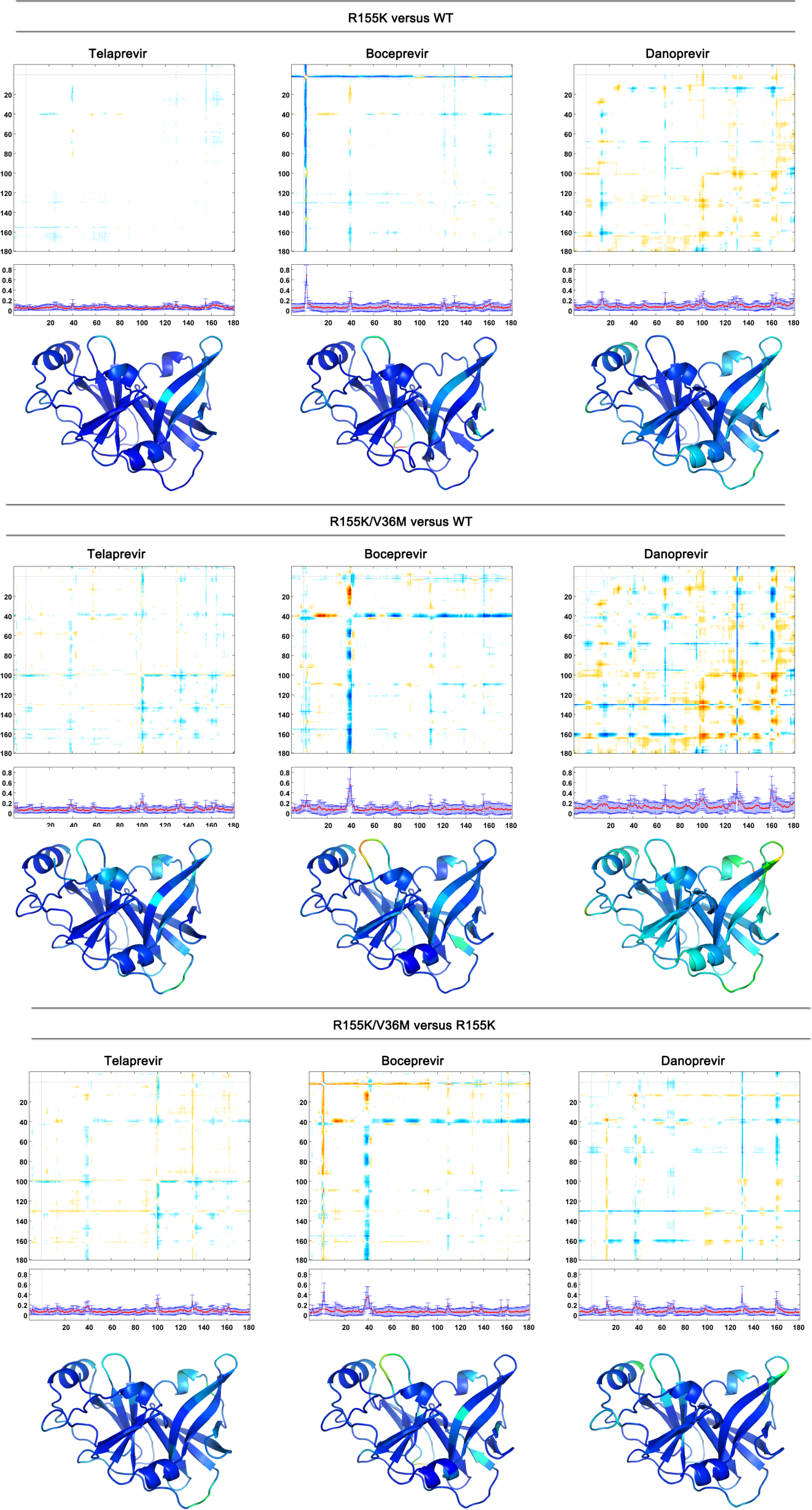
Distance-difference maps showing the backbone perturbations in mutant crystal structures. The core of the protease is relatively unchanged (colored blue on structure) while backbone structural changes occur mostly around the binding site and in the loop regions.

The electrostatic network at the binding site of NS3/4A, involving residues H57–R155– D168–R123, is important for substrate recognition and inhibitor binding (Romano, Laine et al. 2011, Ali, Aydin et al. 2013). Disruption of this network can arise as a mechanism of resistance by weakening inhibitor binding. Therefore, sensitivity of the salt bridges in this network to resistance mutations was assessed (Table 2). R155 can make two salt bridges, one with D81 and the other with D168 in all PI-bound structures. In the wild-type protease simulations, R155–D168 salt bridge is more stable than the R155–D81 salt bridge, consistent with the distances observed in the crystal structures (6.4 Å for D81–R155 and 4.8 Å for R155–D168). While danoprevir binding further stabilizes R155–D168, the linear inhibitors, telaprevir and boceprevir, optimize the electrostatic network on the binding surface such that both the salt bridges exist more than ∼80% of the time. However, R155K mutation favors D81–K155 over K155–D168 in telaprevir and danoprevir complexes but not in boceprevir. Relative stability of K155–D168 in boceprevir complex is dependent on the stability of the H57–D81 salt bridge, since D81 is no longer available for a salt bridge with K155. In boceprevir-bound protease, disruption of the electrostatic network along the binding surface is predominantly orchestrated by R155K mutation as D81–R/K155 salt bridge likelihood drops from 85% in wild-type complex to less than 5% in both R155K and R155K/V36M variants. A slight destabilization of D81–K155 is observed in the telaprevir-bound and danoprevir-bound R155K/V36M proteases (the percentages that D81–K155 salt bridge existed dropped from 89% to 83% and from 84% to 78%, respectively). The last residue in the network, R123, is solvent exposed even in the bound state; therefore the sidechain conformation may be sensitive to crystal contacts also impacting the stability of the other salt bridges in the network. To test this possibility, all the potential salt bridges on the protein surface were evaluated (Table 2). Neither inhibitor binding nor mutations in the protease significantly changed the surface salt bridges, suggesting that the changes observed on the electrostatic network involving H57–D81–R/K155–D168–R123 is due to the mutations in the protease but not alternate conformations of R123 favored in different crystals. Thus, the fluctuations in the salt bridges, combined with the relative susceptibilities of PIs to the R155K mutation, suggest that the electrostatic network along the binding site is a key factor stabilizing the interactions of the bound inhibitor.

### V36M Causes Distal Perturbations in Protease Backbone Conformation and Flexibility

Backbone structural changes in the crystal structures were assessed by computing the differences in the internal Cα-Cα distances between the resistant and wild-type variants bound to each inhibitor (Figure 7). This analysis enables determining the overall conformational perturbation in the protein structure due to resistance mutations without any possible superposition bias (see Materials and Methods).

**Figure 7.**
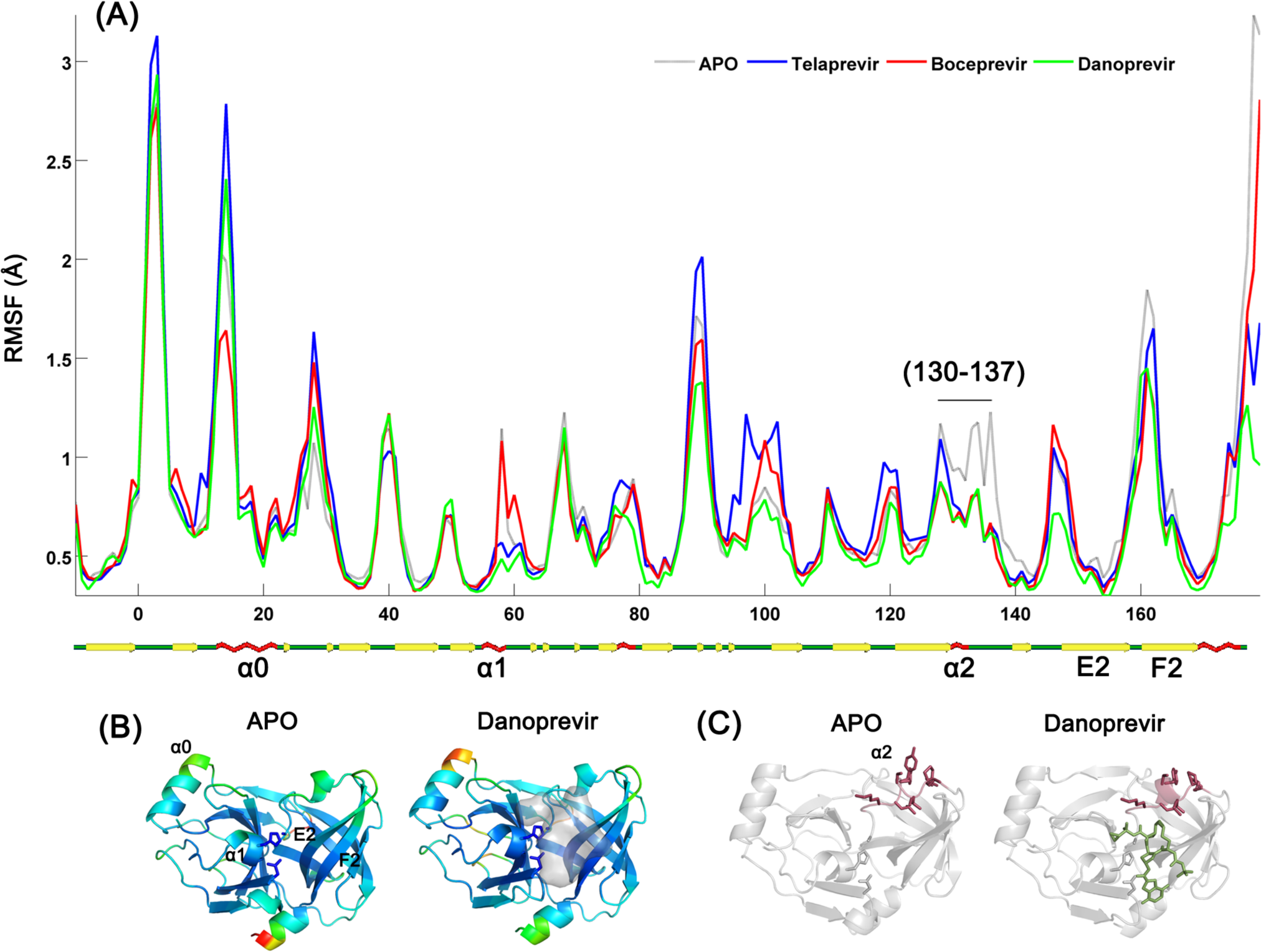
The backbone flexibility of wild-type HCV NS3/4A protease during MD simulations. (A) RMSF values for the protease in the apo form (gray) and in complex with telaprevir (blue), boceprevir (red), and danoprevir (green). (B) The RMSF values for the apo and danoprevir-bound protease are mapped onto the respective crystal structures. (C) The α2 helix (130-137), which is conformationally stabilized upon inhibitor binding, is shown on the crystal structures.

When the single R155K mutation is introduced to the protease, the backbone conformation is the most affected in the macrocyclic danprevir-bound structure. Introducing V36M mutation to the R155K variant further alters the backbone conformation relative to wild-type complex. Backbone of danoprevir-bound protease, which has a ∼300-fold reduction in inhibitory activity due to R155K/V36M, is altered more than the backbone of linear telaprevir- and boceprevir-bound protease with a per-residue average difference of 0.14 Å (for the linear inhibitors: 0.08 Å for telaprevir, 0.09 Å for boceprevir). The V36M distal mutation shifts the danoprevir-bound protease backbone both at the binding site and distal regions of the structure (Figure 7). Significant binding site changes include the loop connecting the E2-F2 strands and the α2 helix, while subtle changes are observed at α1 helix that includes the catalytic H57. Distal regions with substantial backbone perturbations correspond to the loop connecting the β-strands A1 (residue 36 is located on A1) and B1, the α0 N-terminal region, and the α3 C-terminal region. Most of these regions are also flexible and exhibit high fluctuations in the MD simulations (see below and Figure 8). Correspondence between the crystallographic changes in the distance-difference maps and the relative flexibility in MD simulations suggests that the resistance mutations mainly perturb the backbone conformation of relatively flexible regions.

**Figure 8.**
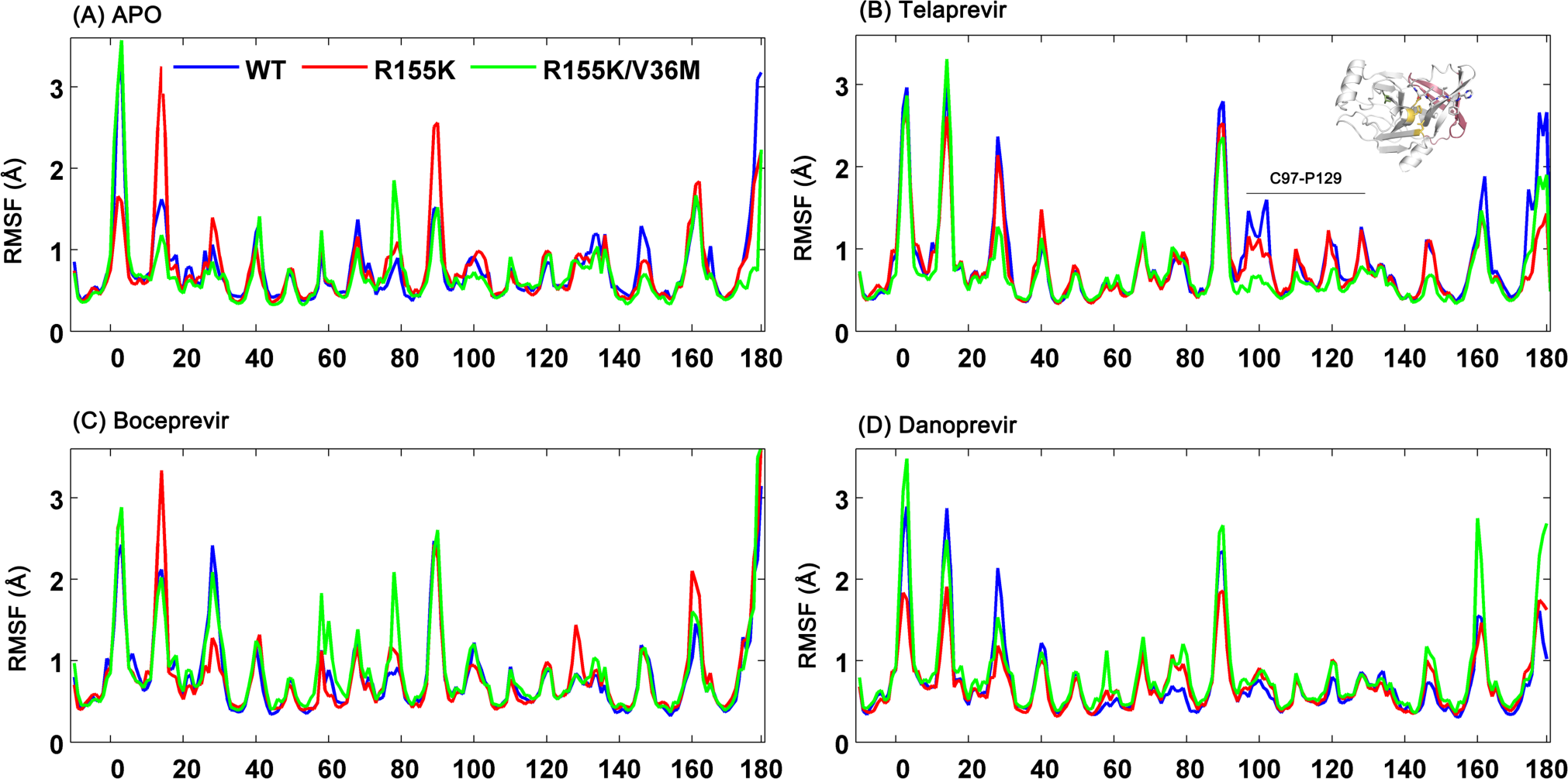
Protease backbone flexibility assessed by atomic positional fluctuations during MD simulations in (A) unbound and (B) telaprevir-, (C) boceprevir-, and (D) danoprevir-bound states of WT (blue), R155K (red), R155K/V36M (green) proteases.

In addition to structural changes in the inhibitor-bound state, resistance mutations can affect the conformational dynamics of the bound and unbound states (Cai, Yilmaz et al. 2012). Overall, the local backbone fluctuations in apo versus inhibitor-bound complexes of wild type NS3/4A protease are comparable (Figure 8). The core of the protein comprised mostly of beta strands is highly stable in all simulations, also supported by the stable secondary structures over the trajectories. Regions with changes in flexibility mainly correspond to loops and helices (with higher RMSF values in Figure 8). The fluctuations of the short α1 helix, which includes the catalytic H57, were suppressed when telaprevir or danoprevir binds but stays apo-like with boceprevir binding. Another region that is in close proximity to the binding site, corresponding to residues 130–137 and including helix α2, lost flexibility when any of the three inhibitors binds (Figure 8), consistent with earlier simulation results (Zhu and Briggs 2011). In this segment of NS3/4A, I132, L135, K136, and G137 and to an extent S138 make favorable vdW interactions with the inhibitor while the backbone amide of G137 hydrogen bonds to the inhibitors. Interestingly, allosteric inhibitors of NS2B/NS3 (Yildiz, Ghosh et al. 2013), another trypsin-like serine protease from dengue virus, bind a region of the enzyme that is topologically similar to the 130–137 segment of HCV NS3/4A. Structural ordering achieved by tight, favorable interactions with the inhibitor implies a role of this 3-10 helix in stabilizing the active site.

We next analyzed changes in backbone fluctuations due to resistance mutations (Figure 9). Relatively restrained residues in the wild-type protease, bound or unbound, are not majorly affected by the resistance mutations. The most striking change in atomic fluctuations with V36M mutation is observed at telaprevir-bound protease’s A2, B2, and C2 strands, which became significantly more restrained compared to the wild-type or R155K single mutant. When bound to danoprevir, residues 70–80, which form E1 strand, were relatively rigid in the wild-type protease and became flexible in both R155K and R155K/V36M variants. In boceprevir-bound protease, this region became flexible only in the double mutant and not in the single R155K mutant. Overall, the resistance mutations changed backbone flexibility not at or near the site of mutation but at other distal regions, and the extent of this change correlated with fold-changes in inhibitor affinity (Figure 9 and Table 1).

**Figure 9.**
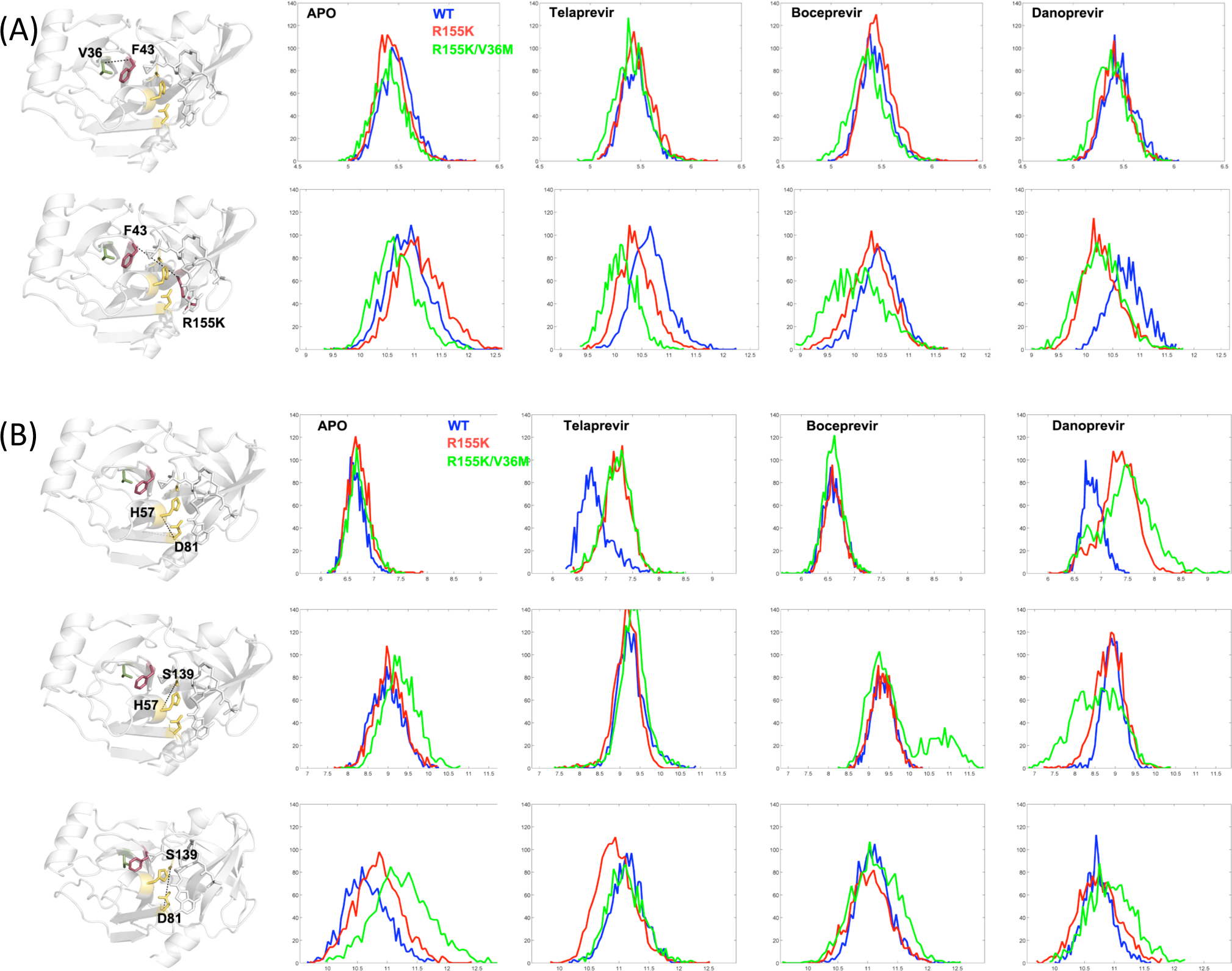
The dynamic distance distribution between protease residue pairs sampled during MD simulations. The distance between (A) F43 and the sites of mutation (V/M36– F43, F43–R/K155), and (B) the catalytic triad residues (H57–D81, H57–S139, D81–S139). The distance (in Å) between the Cα atoms of the two residues is plotted against the percent time that distance was observed.

### The Distal V36M Mutation Changes the Active Site via F43

To better capture the dynamic changes at the binding site due to resistance mutations, inter-residue Cα-Cα distance during MD simulations was tracked for pairs of protease residues. The distance distribution does not considerably change for most of the residue pairs and fluctuate around the distance in crystal structure regardless of mutations. However, V36M mutation changed the dynamics and distance sampled for some key residue pairs across and away from the active site (Figures 10 and S2–S5). In the presence of V36M, residue 36 and F43 become closer to each other compared to WT and R155K complexes (Figure 10A). F43 interacts with the bound inhibitor and is located in B1 strand, which bridges the distal V36M mutation on the A1 strand to the binding site (Figure 1). We found that four residues, K136 in α2 helix and residues 155–157 on E2 β-strand, also became closer to F43 with V36M mutation (Figures 10A and S2). The fact that E2 and B1 β-strands and the α2 helix are closer to each other suggests a slight shrinking in the binding site across the B1–E2 strands direction.

Structural and dynamic reorganization in the binding site involving the α2 helix and B1 and E2 strands also impacted the catalytic triad (Figure 10B). In the apo protease, S139 was pulled away from H57 and D81 with the V36M mutation. In telaprevir and danoprevir complexes, R155K mutation caused H57 and D81 Cα atoms to get farther away from each other, which was further aggravated by V36M in danoprevir-bound protease. When farther apart, the sidechains of these two catalytic residues are not oriented properly for strong salt bridging (Table 2). In contrast, the distances and the H57-D81 salt bridge were conserved in complexes of boceprevir, which is the least susceptible of the three inhibitors to R155K/V36M resistance mutations (Table 1).

Additional residue pairs across the protease indicate changes in the conformational dynamics due to the resistance mutations. The H57-K136 distance decreased in the unbound variants of R155K and R155K/V36M, with a bimodal distribution in R155K/V36M (Figure S4). Since K136 makes key interactions with the natural substrates, one would expect that a change in the relative orientation of K136 with respect to the catalytic H57 could alter the balance in the molecular recognition events in favor of substrate recognition versus inhibitor binding. The loops connecting A1-B1 and E2-F2 strands, which can be thought of as “pseudo-flaps” in analogy to HIV-1 protease, were also closer to each other in the apo and danoprevir-bound proteases while the width of the distribution changed in telaprevir-bound state (Figure S3).

Since the A1 strand, where V36 is located, is in direct contact with the NS4A cofactor in structure (Figure 1), a mutation in A1 may also affect the interactions with NS4A. To assess whether the V36M mutation interferes with the binding of the cofactor NS4A, which aids in the proper folding of NS3, we calculated the distance distributions between the cofactor and NS3 residues as well. Overall, there were no considerable changes except for subtle shifts in the distance between residues V/M36 in A1 and R28 in NS4A in the apo protease, and biomodal distance distribution between E13 and I25 in the inhibitor-bound complexes (Figure S5).

In summary, detailed investigation of co-crystal structures and conformational ensembles coupled with drug susceptibility assays show that the distal site mutation V36M, in the presence of the binding site mutation R155K, alters the binding site shape through changes in conformational dynamics.

## Discussion and Conclusions

Molecular basis of drug resistance in HCV NS3/4A protease conferred by V36M in the R155K background has been investigated using a combination of drug susceptibility assays, co-crystal structures, and MD simulations. Binding assays and replicon studies were consistent in showing that the R155K/V36M double mutant is more resistant to telaprevir, boceprevir, and danoprevir than the single mutant R155K (Table 1). The linear inhibitors are more susceptible to the V36M distal mutation in the presence of R155K than the macrocyclic danoprevir; the fold-change in Ki due to V36M was 9 and 4 for telaprevir and boceprevir, respectively and 2 for danoprevir in the R155K background. However, as danoprevir is highly susceptible to R155K (158-fold change relative to WT), the double mutant had almost 300 times higher inhibition constant than the WT protease.

Why the V36M mutation distal from the active site is selected in combination with R155K and how this mutation is able to impact inhibitor susceptibility is not entirely evident from the crystal structures alone. The impact of the distal V36M mutation on the crystal structures is subtle, with little change in the inhibitor binding mode, intermolecular hydrogen bonds or salt bridges. However, rather than local changes at the binding site, V36M alters the overall protein conformation and dynamics around the active site via changes in interactions with F43 on B1 strand (Figures 7, 10 and S2). Our results suggest that the impact of V36M mutation is propagated to the binding site through B1 β-strand since V36 is located on the A1 β-strand, right next to B1 strand. The fact that F43, on B1 β-strand, is also a resistance mutation site supports this hypothesis. The result of this distal modulation is an effective decrease in the size of the active site. Because the NS3/4A inhibitors typically protrude beyond the dynamic substrate envelope (Ozen, Sherman et al. 2013), this shrinking is expected to impair inhibitor binding more than substrate recognition, and contribute to drug resistance. These results are reminiscent and consistent with the “network hypothesis” and the decrease in active site size we observed previously with HIV-1 protease (Ragland, Nalivaika et al. 2014).

The V36M mutation may also interfere with the binding of the cofactor NS4A, which aids in the proper folding of NS3. Substitution of a valine with a larger sidechain, methionine, alters the local packing. Since the A1 strand containing V36 is in direct contact with NS4A in structure, a mutation in A1 may as well affect the interactions with NS4A. A hindrance in cofactor binding would also interfere with substrate processing efficiency, which may be the reason for the reduced base-level replicative capacity of V36M and R155K/V36M variants of genotype 1b viruses in the absence of drugs (44% and 76% relative to wild-type) (He, King et al. 2008).

The MD simulations here provided additional insights also into the mechanism of resistance due to R155K mutation, which was previously revealed from crystal structures (Romano, Ali et al. 2010, Romano, Ali et al. 2012). The destabilization of danoprevir’s P2 isoindoline in the presence of R155K both in the crystal structures and MD simulations is consistent with earlier simulation results of Pan et al (Pan, Xue et al. 2012), indicating that the severe destabilization of P2 is not a crystallization or force-field dependent artifact. In fact, the P2 moiety samples diverse conformations distinct from the one observed in the crystal structure (Figure 3), with substantial losses in vdW interactions with the active site protease residues.

While the substrate envelope effectively explains the molecular mechanism of resistance to most active site mutations, understanding the effect of mutations distal to the binding site on inhibitor susceptibility is more challenging. Earlier studies on HIV-1 protease showed that drug resistance can be conferred not only via changes in direct interactions with the small-molecule inhibitors and substrates but also alterations in the protein dynamics (Foulkes-Murzycki, Scott et al. 2007, Cai, Yilmaz et al. 2012, Mittal, Cai et al. 2012) and propagated to the active site (Ragland, Nalivaika et al. 2014). Mutations distal to the binding site do not necessarily have an impact on the native structure of the protease-inhibitor complex, however, through changes in conformational dynamics, the exchange rate between conformational states are affected. Consequently, the equilibrium properties of inhibitor binding are negatively impacted. We find that V36M is such a distal mutation, which validates the network hypothesis we previously proposed (Ragland, Nalivaika et al. 2014) and allosterically changes the dynamics at the HCV NS3/4A protease active site to aggravate R155K resistance.

## Materials and Methods

### Mutagenesis and gene information

The HCV genotype 1a NS3/4A protease gene, described in a Bristol-Meyers Squibb patent (Wittekind, Weinheimer et al. 2002), was synthesized by Gen-Script and cloned into the pET28a expression vector (Novagen). Highly soluble NS3/4A protease domain contains 11 core amino acids of NS4A covalently linked at the N terminus. A similar protease construct exhibited catalytic activity comparable to that of the authentic full-length protein (Taremi, Beyer et al. 1998). R155K and R155K/V36M protease variants were generated using the QuikChange Site-Directed Mutagenesis Kit from Stratagene and sequenced by Genewiz.

### Expression and purification

NS3/4A protease expression and purification were carried out as described previously (Gallinari, Brennan et al. 1998, Wittekind, Weinheimer et al. 2002). Transformed Escherichia coli BL21(DE3) cells were grown at 37°C until OD600 reached 0.6 and induced by adding 1 mM IPTG. Cells were harvested after overnight expression at 4°C and pelleted. Cell pellets were resuspended in 5 mL/g of resuspension buffer (50 mM phosphate buffer at pH 7.5, 500 mM NaCl, 10% glycerol, 2 mM β-mercaptoethanol [β-ME]), and lysed with a cell disruptor. The soluble fraction was applied to a nickel column (Qiagen), washed with resuspension buffer supplemented with 20 mM imidazole, and eluted with resuspension buffer supplemented with 200 mM imidazole. The eluant was dialyzed overnight (molecular mass cutoff, 10 kDa) against resuspension buffer to remove the imidazole, thrombin treatment was applied simultaneously to remove the His tag. The nickel-purified protein was then flash-frozen with liquid nitrogen and stored at −80°C for future use.

### Crystallization

Danoprevir was prepared in-house using our convergent reaction sequence as described previously (Romano, Ali et al. 2010); boceprevir was provided by Merck & Co., Inc; telaprevir was purchased from A ChemTek, Inc. (Worcester, MA). For crystallization, the protein solution was thawed and loaded on a HiLoad Superdex75 16/60 column equilibrated with gel filtration buffer (25 mM morpholineethanesulfonic acid [MES] at pH 6.5, 500mMNaCl, 10% glycerol, 30 μM zinc chloride, and 2 mM dithiothreitol [DTT]). The protease fractions were pooled and concentrated to 25 mg/mL using an Amicon Ultra-15 10-kDa device (Millipore). The concentrated samples were incubated 1 h with 2 to 20 molar excess of protease inhibitors. Concentrated protein solutions were then mixed with precipitant solution (20 to 26% polyethylene glycol [PEG] 3350, 0.1 M sodium MES buffer at pH 6.5, and 4% ammonium sulfate) at a 1:1 ratio in 24-well VDX hanging-drop trays and diffraction-quality crystals were obtained overnight.

### Data collection and structure solution

Crystals large enough for data collection were flash-frozen in liquid nitrogen for storage. Constant cryostream was applied when mounting crystal, and X-ray diffraction data were collected at Advanced Photon Source LS-CAT 21-ID-F and our in-house Rigaku_Saturn 944 X-ray system, respectively. The product complexes diffraction intensities were indexed, integrated, and scaled using the program HKL2000 (Otwinowski and Minor 1997). All structure solutions were generated using simple isomorphous molecular replacement with PHASER (McCoy, Grosse-Kunstleve et al. 2007). The model of viral substrate N-terminal product 5A-5B (3M5O) (Romano, Ali et al. 2010) was used as the starting model for all structure solutions. Initial refinement was carried out in the absence of modeled ligand, which was subsequently built in during later stages of refinement. Upon obtaining the correct molecular replacement solutions, ARP/wARP or Phenix (Adams, Afonine et al. 2010) were applied to improve the phases by building solvent molecules (Morris, Perrakis et al. 2002). Crystallographic refinement was carried out within the CCP4 program suite or PHENIX with iterative rounds of TLS and restrained refinement until convergence was achieved (Collaborative-Computational-Project 1994). The final structures were evaluated with MolProbity (Davis, Leaver-Fay et al. 2007) prior to deposition in the Protein Data Bank. Five percent of the data was reserved for the free R-value calculation to prevent model bias throughout the refinement process (Brunger 1992). Manual model building and electron density viewing were carried out using the program COOT (Emsley and Cowtan 2004).

### Drug susceptibility and enzyme inhibition

For enzyme inhibition experiments, 5 nM of the genotype 1a HCV NS3/4A protease domain was incubated with increasing boceprevir concentrations for 90 min in 50 mM Tris assay buffer (5% glycerol, 5 mM TCEP, 6 mM LDAO and 4% DMSO, pH 7.5). Proteolysis reactions were initiated by adding 100 nM HCV NS3/4A substrate [Ac-DE-Dap(QXL520)-EE-Abu-y-[COO]AS-C(5-FAMsp)-NH2] (AnaSpec) and monitored using the EnVision plate reader (Perkin Elmer) at excitation and emission wavelengths of 485 nm and 530 nm, respectively. The initial cleavage velocities were determined from sections of the progress curves corresponding to less than 15% substrate cleavage. Apparent inhibition constants (Ki) were obtained by nonlinear regression fitting to the Morrison equation of initial velocity versus inhibitor concentration using Prism 5 (GraphPad Software). Data were collected in triplicate and processed independently to calculate the average inhibition constant and standard deviation.

### Distance-difference maps

The pair-wise atomic distances between each Cα of a given protease molecule and every other Cα in the same molecule were calculated. The differences of these Cα-Cα distances between each pair of protease molecules were then calculated and contoured as a map for visualization. These maps allow for effective structural comparisons without the biases associated with superimpositions.

### Hydrogen bonds

A hydrogen bond was defined based on the criteria of donor-acceptor distance of less than 3.5 Å and hydrogen-donor-acceptor angle less than 30° and calculated with VMD for both MD trajectories and crystal structures (Humphrey, Dalke et al. 1996). However, the exact coordinates for hydrogen atoms, even in structures of resolution as high as 1.5 Å, cannot be determined with confidence based on the electron density maps. Therefore, for proper evaluation of potential hydrogen bonds in a crystal structure, hydrogen atoms were added to the crystal structures such that the hydrogen bonding network was optimized using Maestro (Maestro v9.2. Portland 2011). An energy minimization was also performed on the crystal structures by completely constraining the non-hydrogen atoms. These structures were then evaluated for the hydrogen bonds formed between the protease and the inhibitors.

### Salt bridges

Salt bridges were defined as an interaction between a side-chain oxygen atom of Asp or Glu within 4.0 Å of a nitrogen atom of Arg or Lys. The salt bridges in the MD trajectories were tracked using VMD (Humphrey, Dalke et al. 1996).

### van der Waals interactions

The protease-inhibitor van der Waals contact energies were calculated by a simplified Lennard-Jones potential, as previously described in detail (Ozen, Haliloglu et al. 2011). Using this simplified potential for each nonbonded protease-inhibitor pair, ∑V(rij) was then computed for each protease and inhibitor residue. The Lennard-Jones parameters, ε and σ, were taken from the OPLS2005 forcefield. For the pairs involving two separate atom types, the parameters were geometrically averaged.

### Molecular dynamics simulations

The crystallographic waters within 4.0 Å of any protein or inhibitor atom were kept; all buffer salts were removed from the coordinate files. To get a more realistic model of the active enzyme interactions with the inhibitors, the A139S back-mutation was introduced in silico to the crystal structures that originally have S139A mutation in the binding site by deleting the Ala sidechain and predicting the conformation of the Ser sidechain using the software Prime (Suite 2011: Prime 3.0 Schrödinger). Because the N-terminal residues in the NS3/4A construct, GSHMASMKKK, were extremely flexible in earlier simulations and did not form stable secondary structure, this region was deleted from the coordinate file to reduce the computational cost (Ozen, Sherman et al. 2013). The structures were prepared for the MD simulations using the Protein Preparation Wizard from Schrödinger (2011, Madhavi Sastry, Adzhigirey et al. 2013). This process adds hydrogen atoms, builds sidechains missing atoms, and determines the optimal protonation states for ionizable sidechains and ligand groups. In addition, the hydrogen bonding network was optimized by flipping the terminal chi angle of Asn, Gln, and His residues and sampling hydroxyl/thiol hydrogens. The exhaustive sampling option with the inclusion of water orientational sampling was used. Following this step, the structure was minimized in vacuum with restraints on heavy atoms using he Impact Refinement module with the OPLS2005 force field and terminated when the root-mean square deviation (RMSD) reached a maximum cutoff of 0.3 Å. This step allows hydrogen atoms to be freely minimized, while allowing heavy-atom movement to relax strained bonds, angles, and clashes.

Desmond(Bowers, Chow et al. 2006) with OPLS2005 force field (Jorgensen, Chandrasekhar et al. 1983, Shivakumar, Williams et al. 2010) was used in all simulations. The prepared systems were solvated in an orthorhombic solvent box with the SPC water model extending 10 Å beyond the protein in all directions using the System Builder utility. The overall charge of the system was neutralized by adding the appropriate number of counterions (Na+ or Cl-).

Each system was relaxed using a protocol consisting of an initial minimization restraining the solute heavy atoms with a force constant of 1000 kcal mol-1 Å-2 for 10 steps with steepest descent and with LBFSG method up to 2000 total steps with a convergence criterion of 50.0 kcal mol-1 Å-2. The system was further minimized by restraining only the backbone and allowing the free motion of the sidechains. At this stage, the restraint on the backbone was gradually reduced from 1000 to 1.0 kcal mol-1 Å-2 with 5000 steps (250 steepest descent plus 4750 LBFSG) for each value of force constant (1000, 500, 250, 100, 50, 10, 1.0 kcal mol-1 Å-2) and finally an unrestrained energy minimization was performed.

After energy minimization, each system was equilibrated by running a series of short MD steps. First, a 10 ps MD simulation at 10 K was performed with a 50 kcal mol-1 Å-2 restraint on solute heavy atoms and using Berendsen thermostat in the NVT ensemble. MD steps were integrated using a two time-step algorithm, with 1 fs steps for bonded and short-range interactions within the 9 Å cutoff and 3 fs for long-range electrostatic interactions, which were treated with the smooth particle-mesh Ewald (PME) method (Darden, Darrin et al. 1993, Essmann, Perera et al. 1995). Time steps were kept shorter at this first MD stage to reduce numerical issues associated with large initial forces before the system equilibrates. This was followed by another restrained MD simulation for 10 ps at 10 K with a 2 fs inner and 6 fs outer time step in NPT ensemble. The temperature of the system was slowly increased from 10 K to 300 K over 50 ps retaining the restraint on the system and 10 ps MD was performed without the harmonic restraints. Production MD simulations were carried out at 300 K and 1 bar for 100 ns using the NPT ensemble, Nose-Hoover thermostat, and Martyna-Tuckerman-Klein barostat. The long-range electrostatic interactions were computed using a smooth particle mesh Ewald (PME) (Essmann, Perera et al. 1995) approximation with a cutoff radius of 9 Å for the transition between the particle-particle and particle-grid calculations and van der Waals (vdW) interactions were truncated at 9 Å. The coordinates and energies were recorded every 5 ps.

### Molecular Modeling

Crystal structures for apo R155K and R155K/V36M variants were not available. These structures were modeled using two separate templates. First, the inhibitor coordinates were removed from the crystal structures of the danoprevir-bound R155K (PDB ID: 3SU0) and R155K/V36M variants and the resulting structure were relaxed using the energy minimization protocol described above. Second, the mutations were introduced to the crystal structure of the wild-type apo protease and the resulting structures were again minimized with the same protocol. To ensure the equivalence of the independently obtained structural models, the energy-minimized structures were then compared with respect to backbone structural changes, side chain orientations of the mutated amino acids and the spatial neighbors of the mutation sites.

## Acknowledgements

This work was supported by the National Institute of Allergy and Infectious Disease (R01-AI085051 and 2R01AI085051-06). We thank Nese Kurt Yilmaz for editorial assistance.

